# Espalier: Efficient tree reconciliation and ARG reconstruction using maximum agreement forests

**DOI:** 10.1101/2022.01.17.476639

**Authors:** David A. Rasmussen, Fangfang Guo

## Abstract

In the presence of recombination individuals may inherit different regions of their genome from different ancestors, resulting in a mosaic of phylogenetic histories across their genome. Ancestral recombination graphs (ARGs) can capture how phylogenetic relationships vary across the genome due to recombination, but reconstructing ARGs from genomic sequence data is notoriously difficult. Here we present a method for reconciling discordant phylogenetic trees and reconstructing ARGs using maximum agreement forests (MAFs). Given two discordant trees, a MAF identifies a set of topologically concordant subtrees present in both trees. We show how discordant trees can be reconciled through their MAF in a way that retains discordances strongly supported by sequence data while eliminating conflicts likely attributable to phylogenetic noise. We further show how MAFs and our reconciliation approach can be combined to select a path of local trees across the genome that maximizes the likelihood of the genomic sequence data, minimizes discordance between neighboring local trees, and identifies the recombination events necessary to explain remaining discordances to obtain a fully connected ARG. While heuristic, our ARG reconstruction approach is often as accurate as more exact methods while being much more computationally efficient. Moreover, important demographic parameters such as recombination rates can be accurately estimated from reconstructed ARGs. Finally, we apply our approach to plant infecting RNA viruses in the genus *Potyvirus* to demonstrate how true recombination events can be disentangled from phylogenetic noise using our ARG reconstruction methods.

## Introduction

Recombination shuffles genetic material between different genetic backgrounds, creating novel combinations of mutations and new genotypes. Recombination is therefore a major driver of both genotypic diversity and phenotypic novelty. For example, recombination contributes to the emergence of pathogen strains with novel combinations of virulence factors [Zhou et al., 1997], resistance genes and antigenic escape mutations [Simon-Loriere and Holmes, 2011]. At the same time, recombination poses a formidable challenge to phylogenetic inference because individuals may inherit different regions of their genome from different ancestors, leading to a mosaic of phylogenetic relationships across the genome that cannot be captured by any single phylogenetic tree [Hudson et al., 1990, Schierup and Hein, 2000].

Despite the complexity recombination introduces to phylogenetic analysis, patterns of ancestry are often highly correlated across large regions of the genome [Wiuf and Hein, 1999, Rasmussen et al., 2014, Kelleher et al., 2016]. This can easily be seen if we think about genomes as being divided into different intervals separated by breakpoints at which recombination events have occurred. If no recombination has occurred within each interval, the ancestry of individuals within each interval can be captured by a single local tree. Adjacent local trees separated by a single recombination event are expected to be very similar because the topological effect of a single recombination event is analogous to one subtree prune and regraft (SPR) move which prunes a subtree from one local tree and then reattaches that subtree to a different parental lineage [Hein et al., 2004]. Neighboring trees on either side of a recombination breakpoint will therefore have similar topologies outside of the lineages directly impacted by a recombination event. However, there can be considerable phylogenetic uncertainty about local trees reconstructed from empirical sequence data, making it difficult to discern if phylogenetic discordance between trees is attributable to recombination or due to phylogenetic “noise” caused by errors in tree reconstruction.

Ancestral recombination graphs (ARGs) capture these differing but correlated patterns of ancestry across the genome as a connected network of coalescent and recombination events [Hudson et al., 1990, Griffiths and Marjoram, 1997, McVean and Cardin, 2005]. Each local tree is embedded within a larger graph such that the ARG can be thought of as a sequence of local trees for each genomic region separated by breakpoints at which recombination events have occurred. A fully reconstructed ARG therefore not only shows how ancestral relationships vary across the genome, but also what lineages have exchanged genetic material and what genetic material was acquired from which ancestor. Given the full ARG, likelihood-based inference of key demographic and population genetic parameters such as recombination rates also dramatically simplifies [Kuhner et al., 2000].

Accurate and efficient methods for reconstructing ARGs from genomic data have therefore been thought of as a “holy grail” in phylogenetics and population genomics. However, little progress has been made in developing ARG inference methods that are both general and computationally efficient. Approaches based on parsimony that seek to find minimal ARGs with the fewest number of recombination events necessary to explain the sequence data generally assume an infinite sites model and ignore the possibility of recurrent mutations, limiting their applicability to rapidly evolving organisms (Song and Hein [2003], Lyngsø et al. [2005], but see Ignatieva et al. [2020]). State-of-the-art methods like ARGweaver employ clever dynamic programming algorithms to sequentially construct ARGs by “threading” lineages between a series of local trees, but are still limited to dozens of genomes [Rasmussen et al., 2014, Hubisz et al., 2020]. Recently, faster but mostly heuristic algorithms have been proposed to work on much larger datasets, generally by approximating the full ARG as a series of local or marginal trees [Kelleher et al., 2019, Speidel et al., 2019]. This may be useful for genealogy-based demographic inference where interest lies primarily in the temporal distribution of coalescent events, but these approaches do not reconstruct the recombination events connecting lineages across local trees, and so cannot be used to infer properties of the recombination process itself.

The ARG reconstruction method described here is based on first reconstructing local trees and then reconciling differences between them using maximum agreement forests (MAFs). Defined more formally below, a maximum agreement forest (MAF) splits two topologically discordant trees into the smallest possible set of subtrees in which the phylogenetic relationships among taxa in each subtree are concordant (i.e. agree) between trees [Hein et al., 1996, Rodrigues et al., 2007, Whidden and Zeh, 2009]. MAFs are central to our approach as they identify subtrees that agree across local trees and therefore allow us to focus only on discordances between local trees. Moreover, we show how MAFs can form the basis of a reconciliation algorithm that preserves discordances between trees that are strongly supported by sequence data while reconciling differences between (local) trees likely attributable to phylogenetic noise. Finally, MAFs can identify both the number and location of recombinations events necessary to reconcile two discordant trees, a property we take advantage of to connect local trees into a fully reconstructed ARG by adding the required recombination events.

We have implemented our MAF-based tree reconciliation and ARG reconstruction methods in a Python package called Espalier, which refers to the ancient agricultural art of controlling woody plant growth by pruning and attaching branches to a common frame such as a trellis. Espalier thus serves as a guiding metaphor for how our algorithm reconciles discordances between trees by pruning and regrafting subtrees. Below, we first describe the MAF algorithm used by Espalier and then show how MAFs can be used to efficiently reconcile discordant trees and reconstruct ARGs. We also show that ARGs reconstructed by Espalier can be used to accurately infer demographic parameters under the coalescent with recombination [Hudson, 1983, Hudson et al., 1990]. Finally, we show how mosaics of discordant phylogenetic trees can be better visualized using tanglegrams by first using Espalier to disentangle discordances likely due to phylogenetic noise.

## Models and Methods

### Tree reconciliation through MAFs

#### Computing Maximum Agreement Forests

Espalier efficiently reconciles discordant phylogenetic trees and reconstructs ARGs using maximum agreement forests. An agreement forest between a pair of trees is a set of subtrees that are topologically concordant between both trees. Topological concordance can formally be defined in terms of bipartitions. Each branch or edge in a tree defines a bipartition of the tree’s leaf taxa set *X* into two disjoint sets. For example, the highlighted edge in tree *T*_1_ in Figure 1A defines the bipartition *AB*|*CDE*. An edge is said to be concordant between two trees *T*_1_ and *T*_2_ if its leaf set bipartition exists in both trees, otherwise the edge is discordant. Two trees are discordant if any edge results in a discordant bipartition of the leaf taxa.

**Figure 1.**
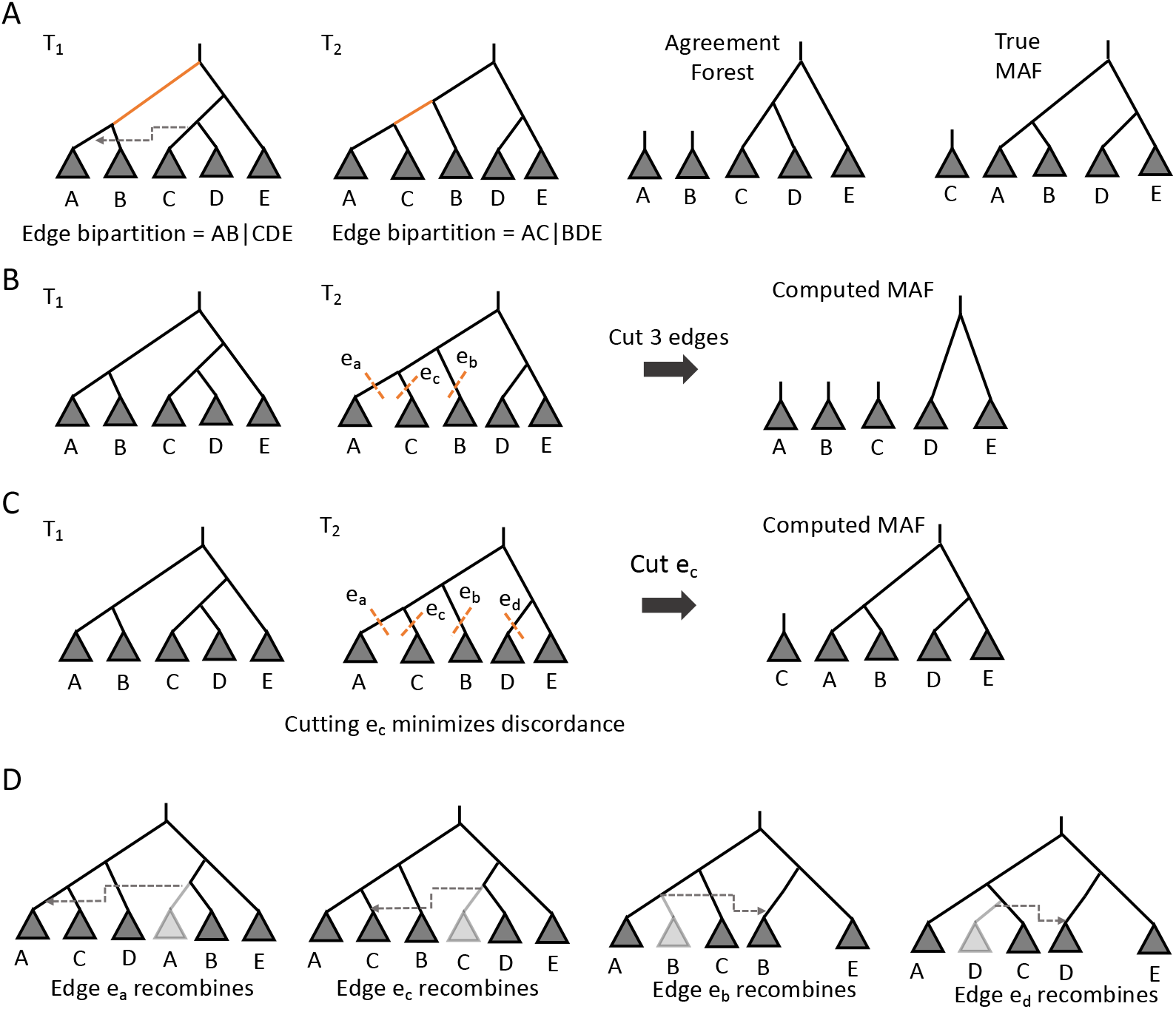
Computing maximum agreement forests. **(A)** A pair of discordant trees *T*_1_ and *T*_2_ where taxa A and B are siblings in *T*_1_ but a recombination event (dashed arrow) moves taxon C such that taxa A and C are siblings in *T*_2_. Also shown is one possible agreement forest with three components, although the true MAF requires cutting only one edge (*e_c_*). Thus the correct SPR distance between the trees is one. **(B)** Schematic illustrating the 3-cut MAF approximation for the same pair of trees. In each step of the algorithm, a discordant edge with children (A,C) in *T*_2_ is chosen and we cut edges *e_a_*, *e_c_* and *e_b_*, where *e_b_* is the sibling of *e_a_* in *T*_1_. Note that cutting one or two of these edges is not always sufficient to reconcile the discordance. For example, if edges *e_a_* and *e_b_* are cut, taxon C would still be sister to D in *T*_1_ but not in *T*_2_. Cutting all three edges guarantees that a discordant edge is removed at the cost of cutting additional edges. **(C)** Schematic illustrating our 4-cut algorithm. The 4-cut algorithm derives from the fact that there are four possible edges surrounding a given discordant edge that may have moved by recombining: edges *e_a_* and *e_c_* in *T*_2_ and their siblings *e_b_* and *e_d_* in *T*_1_. Test cuts are then performed to see which edge should be cut to minimize discordance. In this case, cutting *e_c_* minimizes discordance and returns the true MAF. **(D)** Considering all four possibles edges ensures that, regardless of the the topology of the two starting trees, we make the cut that minimizes discordance. Here for example, if edge *e_c_* recombined, cutting any other edge besides *e_c_* would result in the need to make additional, unnecessary cuts. Note however that all four of these hypothetical edges need not exist. In particular, edges *e_a_* or *e_c_* may not have siblings in *T*_1_, meaning that edges *e_b_* and *e_d_* may not exist (see right two trees).

An agreement forest between two trees can be obtained by cutting all edges such that all component subtrees in the resulting forest are topologically concordant [Rodrigues et al., 2007]. A maximum agreement forest (MAF) is the agreement forest obtained by making the fewest possible cuts and thus has the fewest possible component subtrees. The number of edges that need to be cut in order to obtain the MAF is also the subtree prune and regraft (SPR) distance between two trees [Allen and Steel, 2001], which we denote dSPR. The SPR distance therefore also represents the smallest number of recombination events required to reconcile two topologically discordant trees. An exhaustive search for the MAF would involve cutting all possible combinations of edges and computing the MAF for both rooted and unrooted trees has been shown to be NP-hard [Bordewich and Semple, 2005]. However, Whidden and Zeh [2009] provide a linear time approximation that computes the SPR distance for rooted trees within a factor of three.

The algorithm of Whidden and Zeh [2009] finds, at each step, a discordant edge characterized by having two child nodes that are siblings in *T*_1_ but not in *T*_2_. For example, in Figure 1 the orange edge has child siblings *A* and *B* in *T*_1_ but the sibling of *A* is *C* in *T*_2_. If we let *e_a_* and *e_c_* by the edges leading to the discordant sibling pair (*A, C*) in *T*_2_ and *e_b_* be the edge leading to the sibling of *A* in *T*_1_ (i.e. *B*), cutting all three edges necessarily removes the discordant edge and reduces the dSPR between *T*_1_ and *T*_2_ by one (Figure 1B). Thus, iteratively cutting all three edges surrounding a discordant edge at each step of the algorithm will lead to an approximate MAF with a dSPR within a factor of three of the true dSPR. We therefore refer to this approach as the 3-cut MAF algorithm.

We modify the 3-cut MAF algorithm in order to find the optimal edge to cut at each step in order to avoid making unnecessary cuts. The logic behind our MAF algorithm can be seen by considering all ways in which a discordant edge may arise through a single recombination event or SPR move. Consider a discordant edge that has child siblings (*A, B*) in *T*_1_ but the sibling of *A* is *C* in *T*_2_ as in Figure 1. Depending on the topology of the two trees, this discordant relationship may have arisen because either edge *e_a_* or edge *e_c_* moved through a recombination event (left two trees in Figure 1D) or because a sibling of edge *e_a_* or edge *e_c_*, call these *e_b_* or *e_d_*, recombined and created the sibling pair (*A, C*) in their absence (right two trees in Figure 1D). Thus any one of these four edges (*e_a_*, *e_b_*, *e_c_* or *e_d_*) may have recombined. We therefore introduce a 4-cut algorithm that considers all four possible edges, tests which edge cut minimizes discordance between trees, and then picks the optimal edge to cut (Figure 1C). To ensure the optimal edge is cut we define a distance metric *d_bipart_*(*T*_1_, *T*_2_), which is the number of discordant edges between trees *T*_1_ and *T*_2_, and cut the edge that minimizes *d_bipart_*(*T*_1_, *T*_2_) after the cut is made. Note that in the following algorithm, we assume edges are ordered as in a post-order traversal from tips to root, such that we encounter each discordant edge before we encounter its parent edge.

##### Algorithm 1: The 4-cut MAF approximation

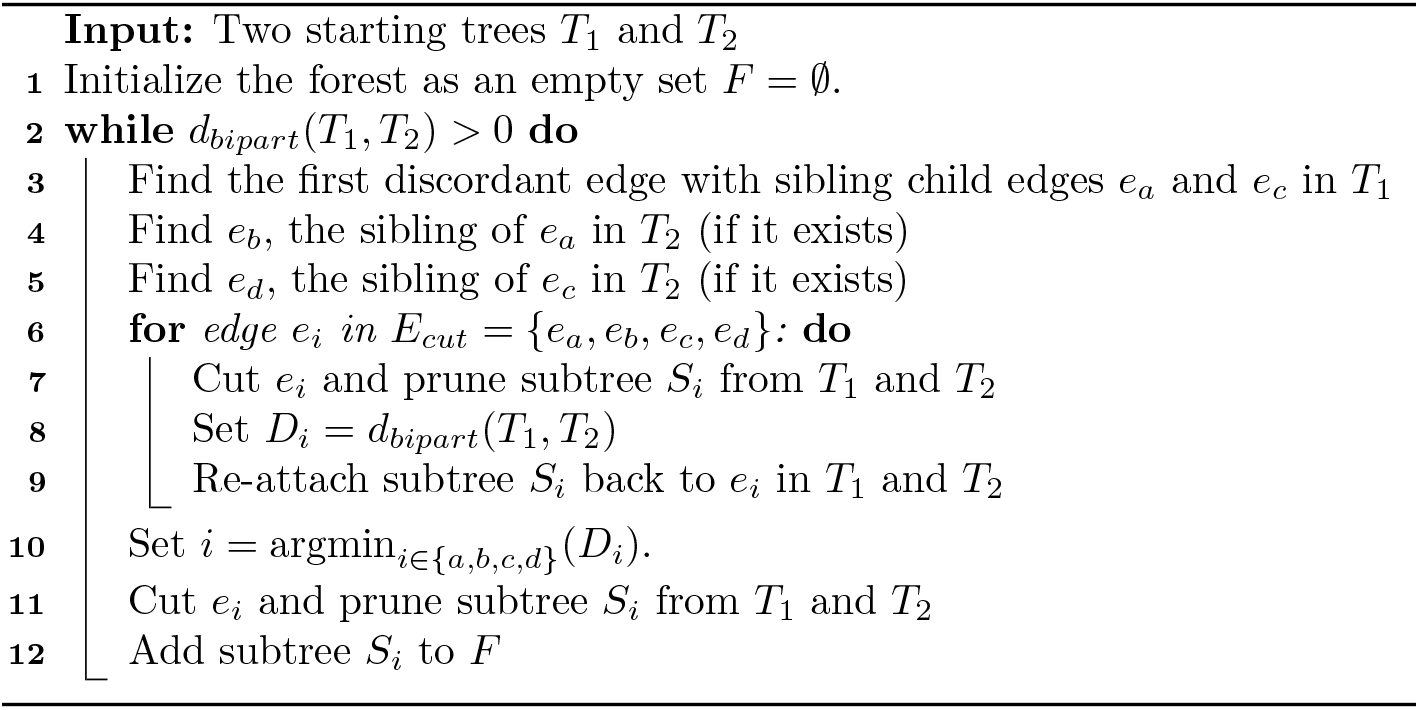

#### Tree reconciliation by iterative regrafting

Next we consider reconciling two discordant trees through their corresponding MAF using an approach we call iterative regrafting. To motivate this approach, consider a local tree *T*_1_ reconstructed from sequence data for a single genomic region and another tree *T*_2_, which could be a local tree for a neighboring genomic region or a reference/consensus tree. The goal is to find a tree that is highly supported by the sequence data over the genomic region being considered while also minimizing discordance between trees by reconciling topological differences that are not strongly supported by the sequence data.

In the iterative regrafting algorithm, we start with a pair of discordant trees *T*_1_ and *T*_2_ and their maximum agreement forest *F*. We will order the subtrees *V* in *F* in the reverse order from which they were pruned by the MAF algorithm, such that *V*_0_ is the last remaining subtree in *F* after all other subtrees were pruned and *V*_1_, *V*_2_, … *V_m_* are ordered such that *V_m_* is the first subtree that was pruned by the MAF algorithm. Starting with *V*_0_, we can regraft *V*_1_ back to its original parent edge from which it was pruned in either *T*_1_ or *T*_2_. If we iteratively regraft subtrees back to their parent edge in the same starting tree, we can reconstruct *T*_1_ and *T*_2_ from the MAF. Note that the ordering of the subtrees is important here because a given subtree may be nested within another subtree, such that if we wanted to regraft a subtree *V*_k_ to its original parent edge, we may need to first regraft another subtree *V_l_* if *k* > *l*. Thus the algorithm reverses the pruning performed in the MAF by regrafting the component subtrees to the starting trees.

We can also reconcile a discordant pair of trees through their MAF by choosing whether each subtree is regrafted back to its parent edge in either *T*_1_ and *T*_2_ creating a reconciled tree *R* that is a topological hybrid of *T*_1_ and *T*_2_ (Figure 2). For each subtree *V_k_* in the MAF, we regraft the subtree to its parent edge in both *T*_1_ and *T*_2_, creating two alternate trees *R*_1_ and *R*_2_. Iterating this procedure leads to a bifurcating algorithm where we regraft the next subtree *V*_*k*+1_ to *R*_1_ along both possible parent edges in *T*_1_ and *T*_2_, and then repeat this regrafting step for *R*_2_. The bifurcating nature of the algorithm therefore generates a search tree, where each internal node represents a partial phylogenetic tree and tip nodes represent full phylogenetic trees. Because two alternative trees are generated at each of the *m* regrafting events, there are 2*^m^* possible reconciled trees.

**Figure 2.**
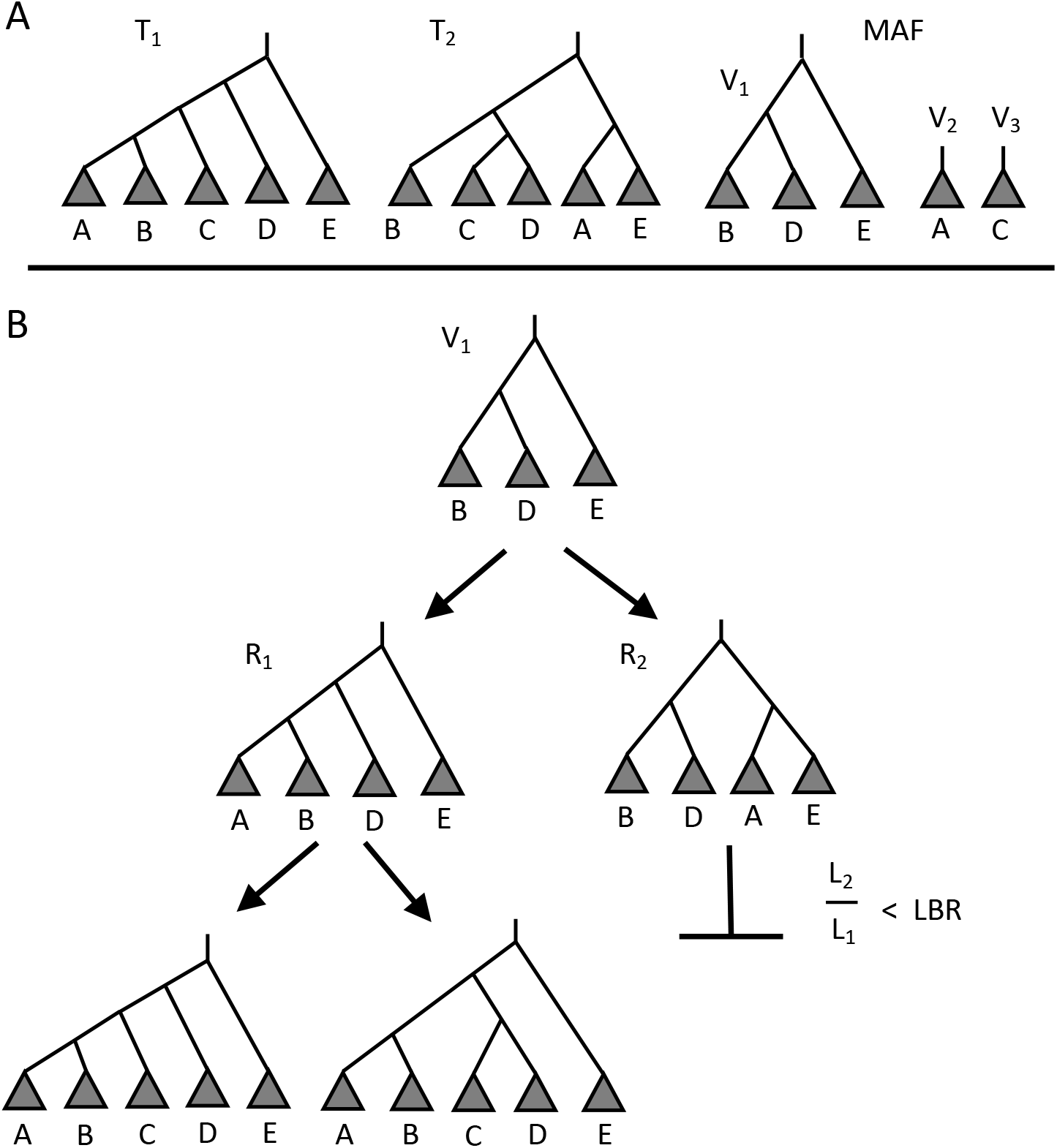
The iterative branch-and-bound regrafting algorithm. **(A)** The algorithm is initialized with a maximum agreement forest (MAF) between a pair of discordant trees *T*_1_ and *T*_2_. **(B)** Starting with the first subtree *V*_1_ in the MAF, the next subtree *V*_2_ is regrafted along both possible parent edges in *T*_1_ and *T*_2_, creating alternative partial trees *R*_1_ and *R*_2_. The likelihoods *L*_1_ and *L*_2_ of the sequence data given *R*_1_ and *R*_2_ are then computed. In this hypothetical scenario, *L*_2_ / *L*_1_ is far less than the lower bound ratio (*LBR*) so the search path descending from *R*_2_ is terminated. The algorithm then returns to *R*_1_ and regrafts the next subtree *V*_3_ to *R*_1_ along both possible edges. The algorithm terminates once all search paths have either terminated or reached a tip node in the search tree, at which point a fully reconciled phylogenetic tree is output.

Because only a small fraction of the 2*^m^* possible reconciled trees will be strongly supported by the sequence data, we employ a branch-and-bound-type algorithm to search for candidate reconciled trees with high likelihoods [Hendy and Penny, 1982]. At each step, we start with a partially reconciled tree *R*, and then regraft the next subtree *V*_*k*+1_ along both possible parent edges *T*_1_ and *T*_2_ creating *R*_1_ and *R*_2_. We then compute the likelihoods *L*_1_ and *L*_2_ of the sequence data given the partial trees *R*_1_ and *R*_2_, where *L*_1_ = *L_θ_*(*S*(*R*_1_)|*R*_1_), *S*(*R*_1_) refers to the sequence data for the subset of taxa in *R*_1_, and *θ* is the known or previously estimated parameters governing sequence evolution. We then use the ratio of the likelihoods *L*_1_ and *L*_2_ as an approximate lower bound in the branch-and-bound algorithm: if the ratio of min(*L*_1_, *L*_2_) over max(*L*_1_, *L*_2_) is below some critical lower bound ratio (LBR), such that one tree is much more strongly supported by the sequence data, we terminate the search along the path with the lower likelihood.

In the following algorithm, we traverse the search tree by adding candidate nodes to an open queue of nodes to be processed. Each node in the search tree is denoted by a tuple (*R, k*) consisting of a partial or full phylogenetic tree *R* and a MAF index *k* that refers to the last subtree *V_k_* regrafted to *R*.

##### Algorithm 2: Iterative branch-and-bound regrafting

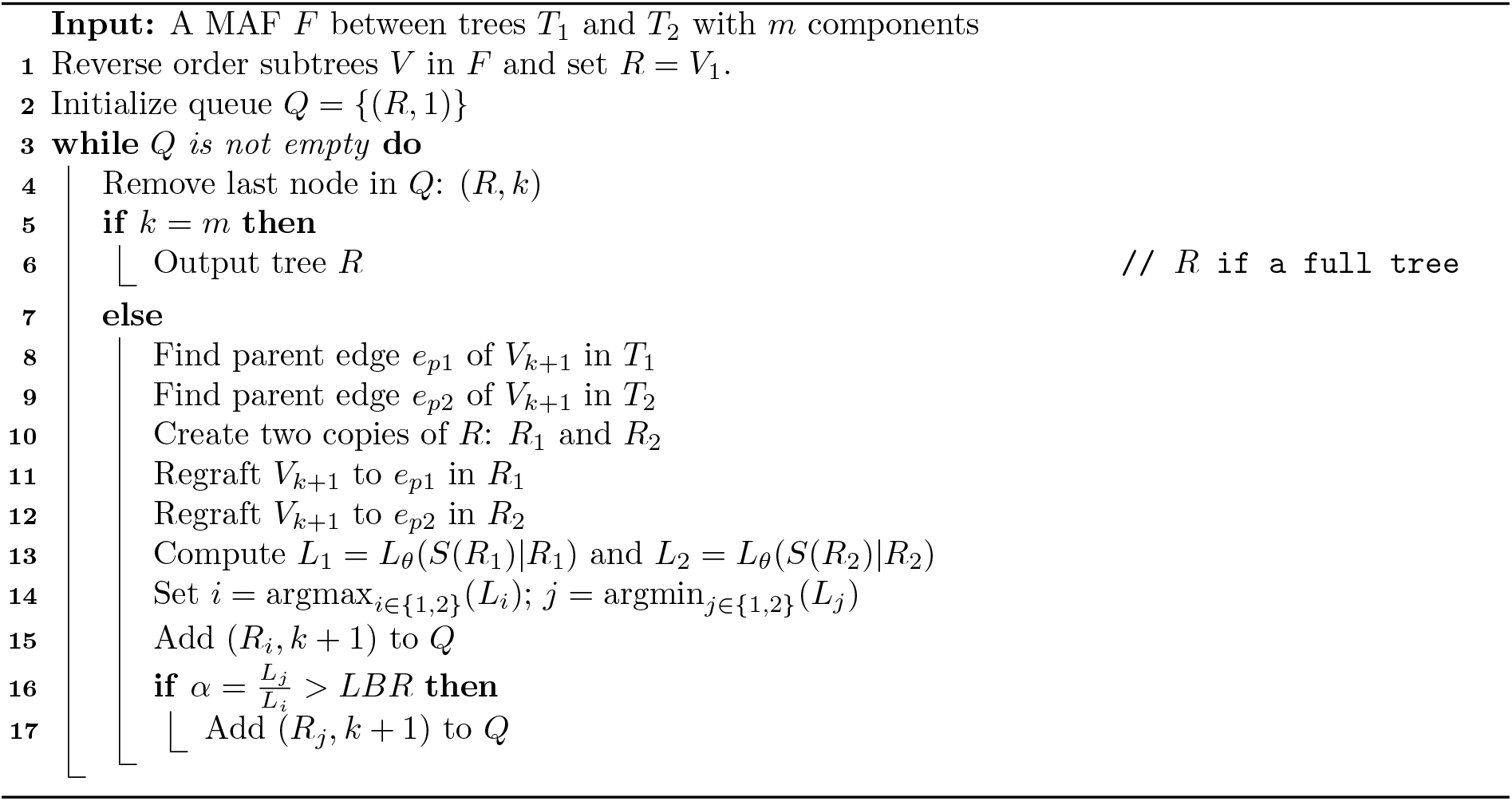

Note that the algorithm is heuristic in that the search for reconciled trees with high likelihood is terminated based on the likelihood ratio of a subset of the sequence data and partial phylogenies. It is therefore possible that terminated search paths would have lead to phylogenies with higher likelihoods *L_θ_*(*S*|*T*) than the trees chosen given the full sequence data. However, in practice we find that this algorithm very efficiently samples trees with high support given the full sequence data.

In the situation where we are trying to reconcile a local tree for a given genomic region against a reference tree, a further improvement to the algorithm is obtained by considering information about the local tree contained in the sequence data from sites neighboring but outside of the genomic region of interest. This is especially true when the genomic region of interest is small such that there are few phylogenetically informative sites within the region. How prior information coming from sites outside the region of interest can be incorporated is described in the Supplemental Methods.

### ARG reconstruction

Espalier efficiently reconstructs ARGs by sampling a sequence of local trees for each genomic region and proceeds in four steps. First, starting trees are reconstructed for each genomic region *i* spanning the closed interval [*l_i_*, *r_i_*], where *l_i_* and *r_i_* refer to the left and right-most genomic positions in the region. Generally, these starting trees will be maximum likelihood (ML) trees inferred from sequence data from the corresponding genomic region *S_i_* ≔ *S_l_i_:r_i__*, although a reference/consensus tree can be substituted if there are not enough phylogenetically informative sites within a genomic region to reconstruct a tree. Second, a set of candidate trees are sampled for each genome region using the tree reconciliation approach described above in order to remove topological differences between local trees that are not well supported by the sequence data. In the third step, described next, we select a sequence of trees, or *tree path*, to include in the ARG. Finally, a full ARG including the recombination events required to explain discordances between neighboring local trees is assembled by imputing plausible recombination events using the MAF algorithm.

#### The tree path sampler

We use a dynamic programming approach based on the Viterbi algorithm [Viterbi, 1967, Forney, 1973] to sample a sequence of local trees, or *tree path*, to form the basis of fully reconstructed ARG. We formulate this problem as a Hidden Markov Model (HMM) model where our objective is to sample a sequence of trees from the joint distribution of local trees *P*(*T*_1:*n*_|*S*_1:*n*_, *r*) over all *n* regions, where:

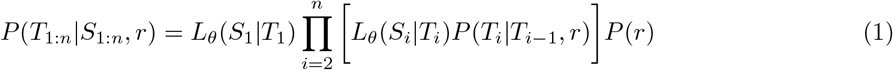

We can see from (1) that the joint distribution can be factored into a product of likelihoods *L_θ_*(*S_i_*|*T_i_*) for the sequence data given the local trees and a product of transition densities *P*(*T_i_*|*T*_*i*–1_, *r*) between local trees, where *r* is the recombination rate per site along each lineage in a tree.

The transition densities provide the probability of moving from tree *T*_*i*–1_ to tree *T_i_* between genomic regions. These transition densities imply that the coalescent-recombination process is Markovian in that the distribution over local tree *T_i_* depends only on the previous local tree *T*_*i*–1_. While this is not necessarily true under the full coalescent with recombination where any two ancestral lineages may recombine [Hudson, 1983], we adopt the Sequential Markov Coalescent (SMC) assumption that ancestral lineages that share no overlapping genetic material are not allowed to recombine [McVean and Cardin, 2005]. In essence, the SMC places additional restraints on the effect of a recombination event in terms of the SPR move it induces: subtrees pruned by recombination can only re-attach to lineages present in the previous local tree rather than any lineage in the entire ARG. While approximate, the SMC is known to be an excellent approximation to the full coalescent with recombination [Wilton et al., 2015].

Here we set the transition probabilities equal to the probability of the minimum number of recombination events necessary to explain the discordance between trees, which is equivalent to the subtree prune and regraft distance *d_SPR_*(*T_i_*, *T*_*i*–1_) computed by the 4-cut MAF algorithm. If recombination events occur independently at rate λ*_rec_* over a genomic region, then the number of recombination events follows a Poisson distribution and the transition density between local trees becomes:

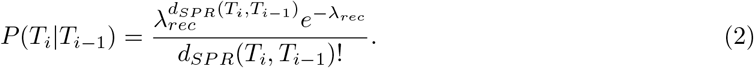

The total rate λ*_rec_* at which recombination events occur within a given region *i* depends on the recombination rate per site *r*, the number of sites at which recombination could have occurred *b_i_* = *l_i_* – *r_i_*, and the total time elapsed along all lineages in the tree, which is the total length of the tree *L_T_i__*. The total recombination rate is therefore:

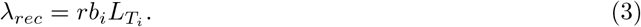

An additional subtlety arises because the SPR distance only considers recombination events that result in an observable change in tree topology. However, certain types of recombination events can occur which do not alter the tree topology such as when the two parent lineages of a recombinant lineage coalesce with one another before coalescing with another lineage in the tree [Hudson and Kaplan, 1985, Hein et al., 2004]. Topology-changing recombination events will therefore occur at a lower rate than λ*_rec_*, resulting in a *thinned* Poisson process [Deng et al., 2020]. Our overall recombination rate therefore needs be reduced by a factor proportional to the probability of a topology-changing recombination event. Deng et al. [2020] provide a formula for the probability of a recombination event not changing the tree topology *P*(*unchanged*). We can therefore rescale λ*_rec_* by 1 – *P*(*unchanged*) to adjust for the rate at which topology changing recombination events occur.

For efficiency, we constrain the state space of *P*(*T_i:n_*|*S*_1:*n*_, *r*) to only include the *reconciled* trees sampled for region *i* using the iterative regrafting algorithm. These candidate trees are placed in a *trellis* 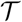 of trees which may contain a variable number of trees for each genomic region. We then use a Viterbi algorithm to select the most likely sequence of trees 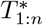 from *P*(*T*_1:*n*_|*S*_1:*n*_, *r*). The algorithm consists of a forward pass in which path probabilities are computed conditional on the sequence data observed up to region *i* and then a backwards pass in which the most likely path is selected given all observed sequence data.

##### Algorithm 3: The Viterbi tree path sampler

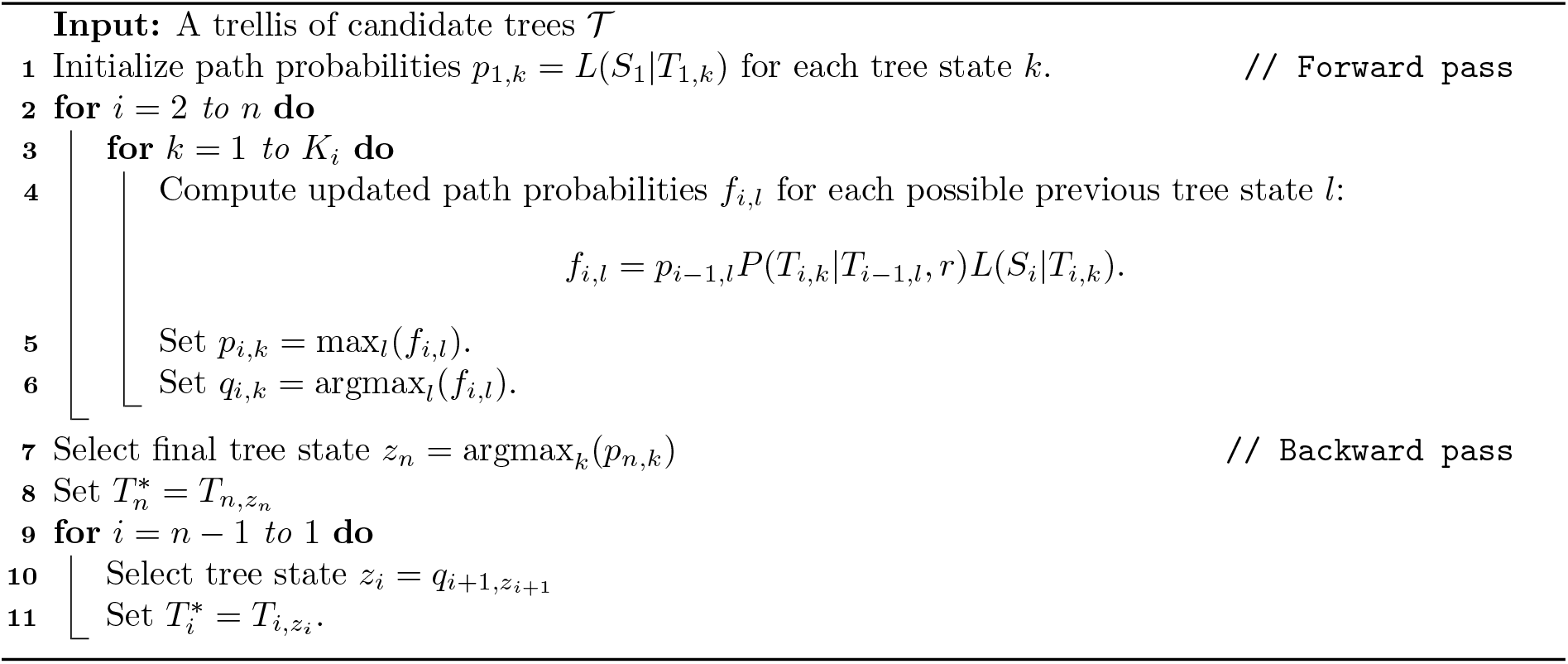

The algorithm returns the maximum a posteriori (MAP) tree path, which maximizes the likelihood of the sequence data while the transition probabilities constrain the path to be consistent with a given recombination rate. In particular, if the the recombination rate is low, the algorithm has a natural “smoothing” effect on the selected tree path in that the transition densities enforce the requirement that neighboring local trees are topologically similar. Finally, while the Viterbi algorithm finds the single most likely path, we describe an alternative forward-backward algorithm in the Supplemental Methods that allows us to randomly sample tree paths with probability proportional to the joint distribution over trees *P*(*T*_1:*n*_|*S*_1:*n*_, *r*). While not used here, this algorithm might be useful to sample a set of highly probable tree paths consistent with the data in order to quantify uncertainty in the local trees or may be used in a MCMC algorithm where we wish to sample tree paths from their posterior distribution conditional on the current value of *r* or other parameters.

#### Rules for sampling recombination events

After sampling a tree path, we still need to sample a set of plausible recombination events to reconcile the topological discordances between neighboring trees and connect recombining lineages in the ARG. For this task, we again use the MAF computed between discordant neighboring trees to identify which lineages likely recombined. Because each component of a MAF represents a subtree that attaches to different edges in *T*_1_ and *T*_2_, identifying the edges to which a subtree attaches in *T*_1_ and *T*_2_ also identifies the lineages that likely recombined. More precisely, if *e_r_* is a cut edge leading to a subtree in the MAF and *e_r_* attaches to edges *e*_*a*1_ and *e*_*a*2_ in trees *T*_1_ or *T*_2_, respectively, then we call *e_r_* the recombinant edge and the attachment edges *e*_*a*1_ and *e*_*a*2_ the parent edges (Figure 3).

**Figure 3.**
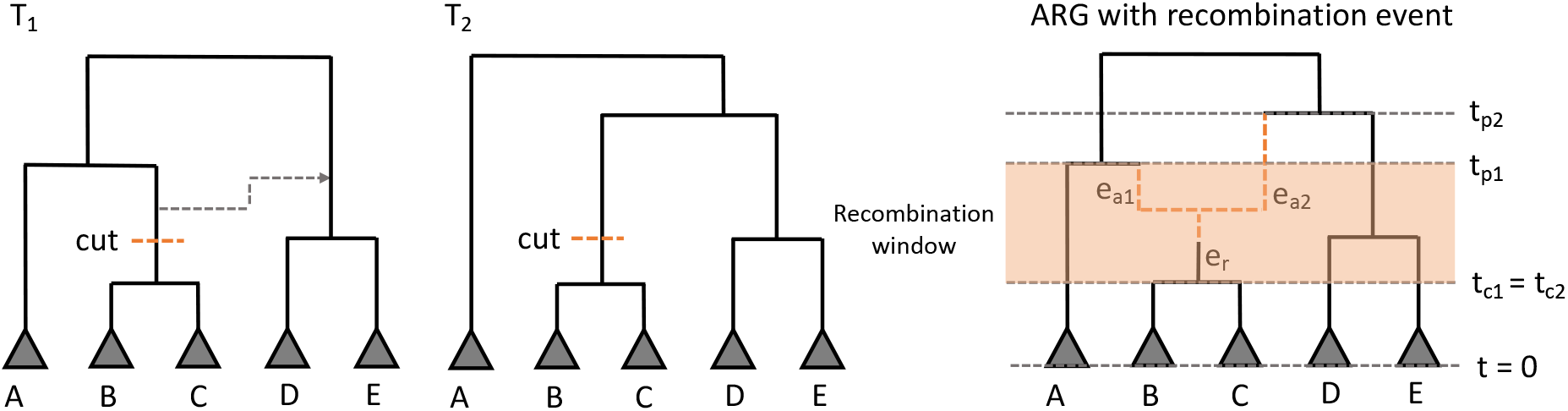
Determining plausible recombination events and times from a MAF. Trees *T*_1_ and *T*_2_ differ due to a recombination event (dashed arrow) in which (B,C) moves. A MAF between *T*_1_ and *T*_2_ is therefore obtained by cutting (B,C) from both trees. Edge *e_r_* leading to (B,C) is therefore the most likely recombinant lineage in the ARG with *e*_*a*1_ and *e*_*a*2_ the most likely parents since *e_r_* attaches to these lineages in *T*_1_ and *T*_2_. Although we do not know the exact timing of the recombination event, the height of the nodes surrounding the recombinant lineage constrains the event to occur within a defined recombination window (shaded interval). If we let *t*_*c*1_ and *t*_*p*1_ be the child and parent node times of the recombinant lineage in *T*_1_ and likewise *t*_*c*2_ and *t*_*p*2_ be the child and parent node times in *T*_2_, then the recombination window is constrained to be between max(*t*_*c*1_, *t*_*c*2_) and min(*t*_*p*1_, *t*_*p*2_), where time increases going backwards from the present at *t* = 0. Note in this case, *t*_*c*1_ = *t*_*c*2_ but this need not be the case if node heights differ between the two trees.

We can therefore add a recombination event by inserting an additional recombination node along the recombinant edge *e_r_* and connecting this node to its two parent edges *e*_*a*1_ and *e*_*a*2_. However, sequence data will contain no information about the exact timing of recombination events. We therefore assign recombination events random times (i.e. node heights) that satisfy the constraints imposed by the heights of the surrounding nodes. In particular, we enforce the requirement that the recombination event must occur during the interval of time in which the parents *e*_*a*1_ and *e*_*a*2_ are present in both trees and thus eligible to recombine. To specify these exact constraints, let *t*_*p*1_ and *t*_*p*2_ be the height of the parent nodes of the recombinant edge in *T*_1_ or *T*_2_. Further, let *t*_*c*1_ and *t*_*c*2_ be the height of the child nodes of the recombinant edge in *T*_1_ or *T*_2_. These times then define a recombination window in which *e*_*a*1_ and *e*_*a*2_ are present in the trees and we can randomly draw a recombination time *t_r_* within this window that satisfies the constraints:

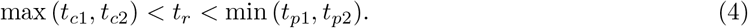

### Demographic inference under the coalescent with recombination

Most ARG reconstruction methods including ours assume that the recombination rate *r* is known, although inferring *r* from empirical data is notoriously difficult [Stumpf and McVean, 2003]. We therefore propose a likehood-based approach to estimate the recombination rate (or other demographic parameters *θ*) together with the ARG that jointly maximize the likelihood of the sequence data using an Expectation-Maximization (EM) algorithm [Dempster et al., 1977]. Note that technically the algorithm is a version of the Baum-Welch algorithm for hidden Markov models, which is a special case of the more general EM algorithm [Fraser, 2008]. In the *expectation step*, we use the Viterbi algorithm to select the tree path *T*_1:*n*_ that maximizes *P*(*T*_1:*n*_|*S*_1:*n*_, *r*) using the current value of the recombination rate *r*. We then impute probable recombination events as described above and add recombination nodes to the local trees to produce a fully connected ARG 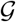. In the *maximization step*, we then find the parameters *θ* that maximize the likelihood of the current ARG by numerical optimization. For our purposes here, the likelihood 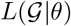 of the ARG is computed under the coalescent with recombination [Hudson, 1983, Hudson et al., 1990], as described in [Kuhner et al., 2000]. Alternately updating the ARG in the expectation step and demographic parameters in the maximization step iteratively until the EM algorithm converges on stable values of **θ** produces a maximum likelihood estimate of both the ARG and *θ*.

#### Algorithm 4: The EM algorithm

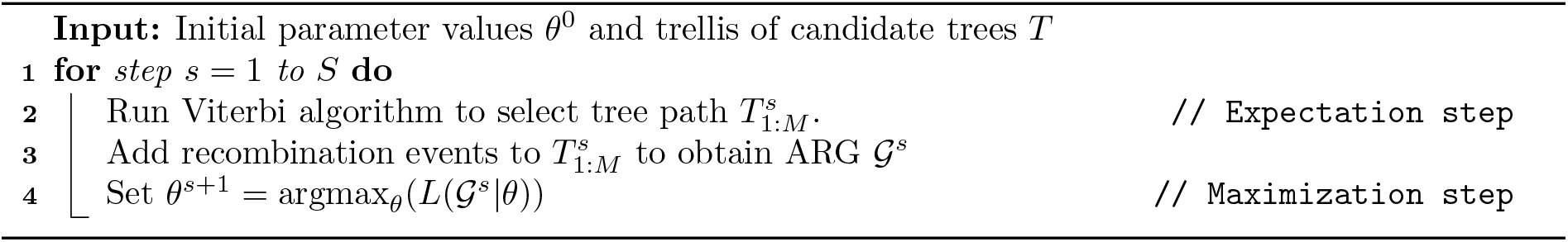

### Implementation of Espalier

The tree reconciliation and ARG reconstruction methods, including the 4-cut MAF algorithm, were implemented in the Python package Espalier. Source code for Espalier is available at github.com/davidrasm/Espalier and documentation is available at espalier.readthedocs.io.

Espalier in turn makes use of several existing phylogenetic packages. Trees are stored and manipulated using Dendropy version 4.5.1 [Sukumaran and Holder, 2010]. We make special use of the way Dendropy encodes edge bipartitions (splits) of the leaf taxa set as integer-valued bitmasks, allowing equivalent edge bipartitions representing the same edge across different trees to be identified through their bitmasks. The bitmask encoding of edge bipartitions also facilitates the rapid identification of topological discordances between trees by comparing the set of edge bipartitions present in each tree using sets of bitmasks.

Full ARGs reconstructed from tree paths are stored and accessed through tree sequence objects in tskit [Kelleher et al., 2016, 2018]. The tree sequence data structure allows ARGs to be stored as flattened tables of nodes and edges. Nodes and edges present across multiple local trees in the ARG therefore only need to be stored once in these tables, allowing for a more concise encoding of local trees in an ARG. Tree sequences also facilitates algorithmic operations on ARGs, such as computing the likelihood of the ARG under the coalescent, as described in Guo and Rasmussen [2022].

Initial maximum likelihood (ML) trees are reconstructed from the sequence data corresponding to each genomic region using RAxML-NG [Kozlov et al., 2019]. We also compute the likelihood of the sequence data at any site in an alignment given a tree using the *–sitelh* method in RAxML-NG. For computational efficiency, we only estimate the molecular evolutionary parameters including substitution rates once during the initial tree reconstruction phase, then use the MLE of each parameter when computing the likelihood of sequence data in the tree reconciliation and ARG reconstruction phase.

### Simulation experiments

To test the performance of Espalier in reconstructing ARGs, mock ARGs were simulated using msprime [Kelleher et al., 2016]. In all simulations, the haploid effective population size *N_e_* = 1, such that time is scaled relative to the rate at which pairs of lineages coalesce, but we vary the mutation rate *μ* to explore how different levels of genetic diversity impact ARG reconstruction. The average pairwise genetic diversity between nucleotide sequences in our simulations is therefore π = 2*N_e_μ* = 2*μ*. Nucleotide sequences for sampled lineages were then simulated along each local tree under a HKY substitution model using pyvolve [Spielman and Wilke, 2015].

### Potyvirus ARG reconstrucion

While currently 167 species of potyviruses are officially recognized [Wylie et al., 2017], we limit our analysis to 131 species for which high-quality, full-length genomes are publicly available. Viral genomes were retrieved from NCBI GenBank using the accession numbers provided by [Gadhave et al., 2020]. Full genome nucleotide sequences were aligned using MAFFT version 7 [Katoh and Standley, 2013]. Potential recombination events with significant signal and their corresponding breakpoint locations were identified using 3SEQ [Lam et al., 2018, Boni et al., 2007]. Because a large number of potential breakpoints were identified across the entire genome including several within each protein coding region, we partitioned the full alignment into non-recombinant blocks (NRBs) within which no recombination breakpoints were identified (see Figure 8A). The 10 longest NRBs with lengths greater than 200bp were selected for further phylogenetic analysis. ML phylogenetic trees for each NRB were reconstructed in RAxML-NG. The Ryegrass mosaic virus reference genome (GenBank accession Y09854.1) was used as an outgroup to consistently root each ML trees. Ryegrass mosaic virus was chosen as it is the type species of the genus *Rymovirus* which recent phylogenetic analyses place sister to the potyviruses [Gibbs et al., 2020]. The resulting ML trees were then time-calibrated using least squares dating in LSD version 0.3 [To et al., 2016] to get an ultrametric tree with all contemporaneously sampled potyviruses equidistant from the root. Due to the difficulty of estimating a consistent molecular clock rate from contemporaneously sampled potyviruses [Gibbs et al., 2020], the molecular clock was fixed at 1.0 substitutions per unit time such that branch lengths in the dated trees are proportional to substitutions per site. Finally, the recombination rate and a reconstructed ARG were inferred using the EM algorithm in Espalier based on the same 10 NRBs.

Tanglegrams were then used to visualize how phylogenetic relationships vary between local ML trees and local trees in the ARG. Tanglegrams were plotted using the baltic package https://github.com/evogytis/baltic in Python.

## Results

### Testing the MAF algorithm

We first tested the MAF algorithm by simulating a random coalescent tree and rearranging its topology through a randomly chosen number of SPR moves. We then used both the 3-cut algorithm of Whidden and Zeh [2009] and our 4-cut algorithm to compute the MAF between the original and randomized trees. Ideally, the computed MAF should identify the correct SPR distance between trees and contain the same number of subtree components (excluding the final subtree) as the true number of SPR moves performed. As expected, the 3-cut algorithm generally overestimates the number of components in the MAF and therefore also the SPR distance by a factor of up to three (Figure 4A&C). In contrast, the 4-cut algorithm generally identifies the correct MAF distance (Figure 4C&D). However, for both algorithms, the SPR distance computed from the MAFs is rather variable and may be either higher or lower than the true number of SPR moves when the tree size (number of tips) is small (Figure 4A&B). This is most likely due to the fact that two or more SPR moves may rearrange the same edge or region of the tree twice, reversing the effect of the original SPR or adding an additional rearrangement. In either case, the 4-cut algorithm is no longer guaranteed to find the optimal cut nor the true number of SPR moves. Increasing the tree size reduces the probability of a SPR move rearranging the same tree region more than once, in which case there is far less variability in the computed SPR distances and the 4-cut algorithm almost always estimates a SPR distance close to the true distance (Figure 4C&D).

**Figure 4.**
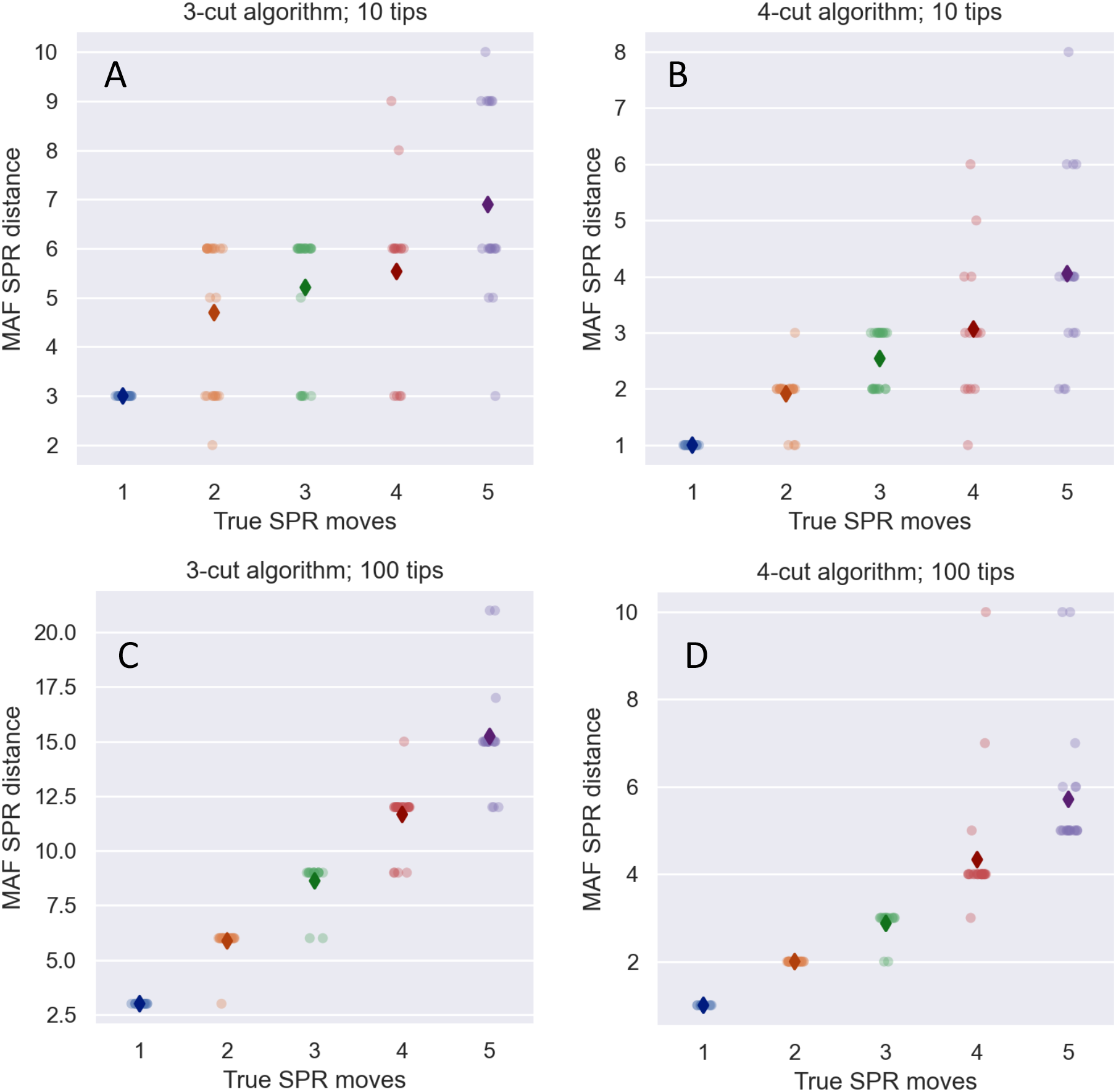
Performance of the 3-cut versus 4-cut approximations in computing maximum agreement forests. In each simulation, a coalescent tree was simulated and then a random number of SPR moves ranging from one to five were performed to topologically rearrange the tree. Performance was tested by comparing the true number of SPR moves in each simulation to the SPR distance (number of subtrees in MAF minus one) identified by either the 3-cut **(A & C)** or the 4-cut algorithm **(B & D)**. Trees included 10 tips in A and B and 100 tips in C and D. 100 simulations were performed under each algorithm and diamonds represent the mean computed SPR distance across simulations for a given number of true SPR moves.

### Testing the reconciliation algorithm

To test our reconciliation approach, we simulated a pair of local trees *T*_1_ and *T*_2_ for neighboring genomic regions in a hypothetical ARG. The simulations were conditioned on exactly one topology-altering recombination event occurring between regions, such that the true SPR distance between trees is always one. A maximum-likelihood local tree, 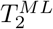, was then reconstructed from simulated sequence data for region 2. 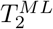 was then reconciled against *T*_1_ (assumed here to be known) using the iterative regrafting algorithm, which by design should resolve discordances between the trees that are not strongly supported by the sequence data. To ensure our algorithm works correctly, we compared normalized Robinson-Fould distances [Robinson and Foulds, 1981] between 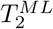 and the true local tree *T*_2_ before and after reconciling to see if reconciliation improves local tree inference. We also compared SPR distances between 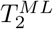 and *T*_1_ before and after reconciliation to check if reconciling local trees correctly removes spurious discordances caused by errors in phylogenetic reconstruction.

We find that reconciling local trees considerably improves local tree reconstruction. After reconciliation, 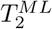 is generally much closer to the true tree *T*_2_ than before reconciliation in terms of RF distances (Figure 5A). The improvement is especially dramatic when the sequence length of region 2 is short such that there is limited phylogenetic information to reconstruct 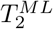. Likewise, after reconciliation the SPR distance between 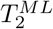 and and *T*_1_ is generally much smaller and closer to the true SPR distance of one (Figure 5B). These two observations suggest that reconciliation correctly eliminates spurious discordances between trees and that reconciling local trees against neighboring trees improves inference of local tree topologies, especially when local trees are reconstructed from limited genomic data.

**Figure 5.**
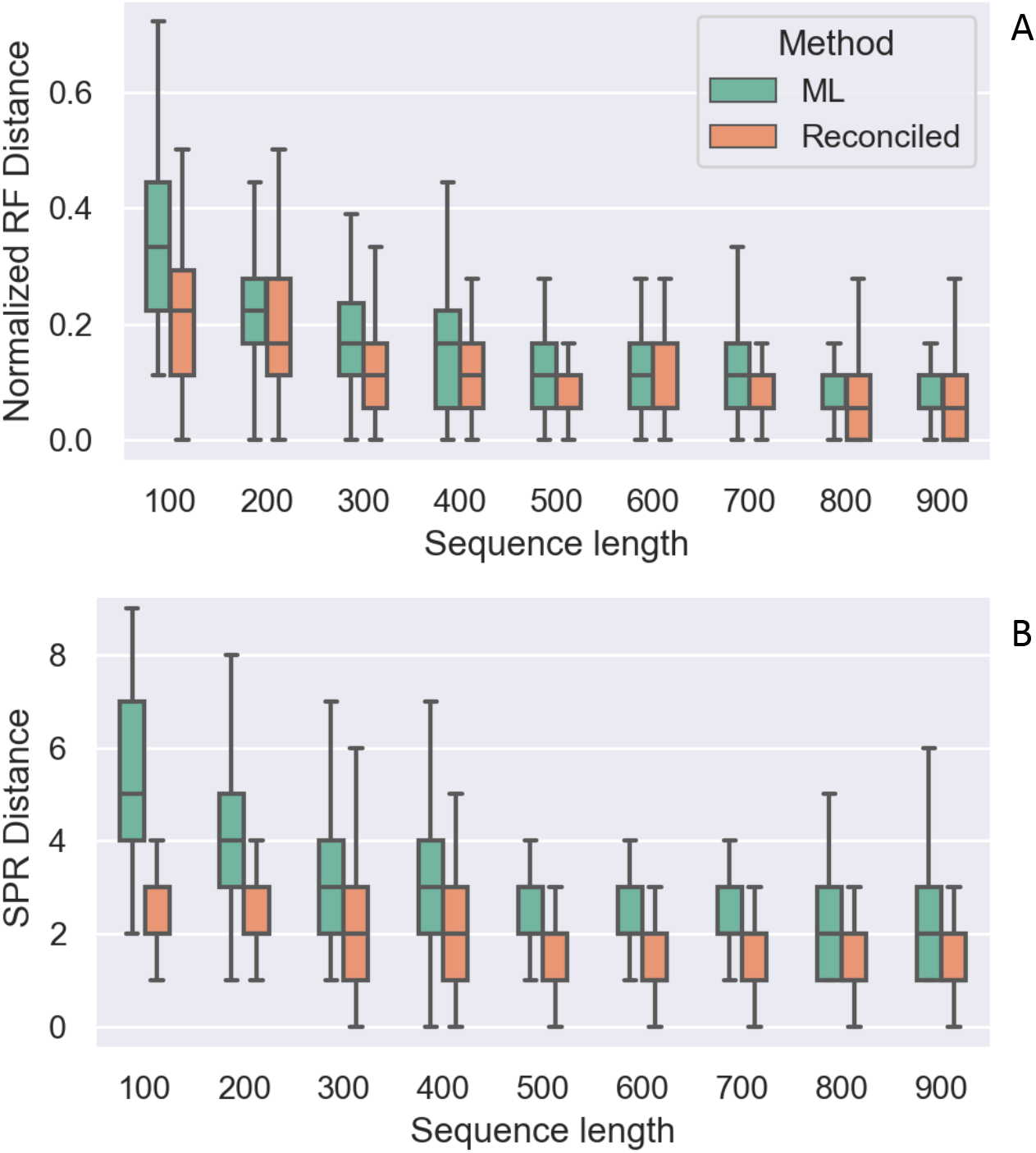
Reconciliation correctly eliminates discordances between trees caused by phylogenetic noise. Two neighboring local trees *T*_1_ and *T*_2_ were simulated but *T*_1_ was assumed known whereas a maximum likelihood tree, 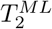, was reconstructed for the second tree using sequence data of varying lengths. Distances between 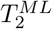 and the true trees were then compared before and after reconciling 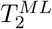 against *T*_1_. **(A)** Normalized Robinson-Fould distances between 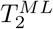 and the true local tree *T*_2_ before and after reconciliation. **(B)** SPR distances between 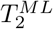 and the true local tree *T*_1_ before and after reconciliation. 100 simulations were performed at each sequence length using local trees with 20 tips. Sequences were simulated under a HKY substitution model with a mutation rate *μ* = 0.1 such that the average pairwise diversity π = 0.2.

### ARG reconstruction performance

Next, we tested the ability of Espalier to accurately reconstruct ARGs from data simulated under the coalescent with recombination. Reasoning that the number of phylogenetically informative sites (SNPs) between recombination breakpoints is likely the key factor limiting our ability to accurately reconstruct ARGs, we fixed the recombination rate *r* = 0.001 per site per generation and varied the mutation rate. To quantify the accuracy of ARG reconstruction, we use normalized Robinson-Fould distances to quantify the topological similarity of local trees in the reconstructed ARG to the local trees in the true (simulated) ARG. We also compare SPR distances between local trees in the reconstructed and true ARGs to check if the reconstruction methods can correctly differentiate true discordance caused by recombination from phylogenetic noise in tree reconstruction. The performance of Espalier was then compared against ARGweaver [Rasmussen et al., 2014], which serves as a gold-standard for comparison since ARGweaver samples ARGs from their exact posterior distribution up to the approximations made by the sequential Markov coalescent model.

At low mutation rates (*μ* = 0.01; *μ*/*r* = 10; π = 2*N_e_μ* = 0.02), the local trees in ARGs reconstructed by both Espalier and ARGweaver are quite topologically distant from the true trees as quantified by the normalized RF distances (Figure 6A), suggesting that both methods introduce considerable phylogenetic error in reconstructing local tree topologies. Nevertheless, the local trees sampled by Espalier and ARGweaver are much closer to the true local trees than maximum likelihood trees reconstructed for each genomic region individually, demonstrating that these methods do correctly leverage information across neighboring genomic regions to reconstruct more accurate local trees (Figure 6A).

**Figure 6.**
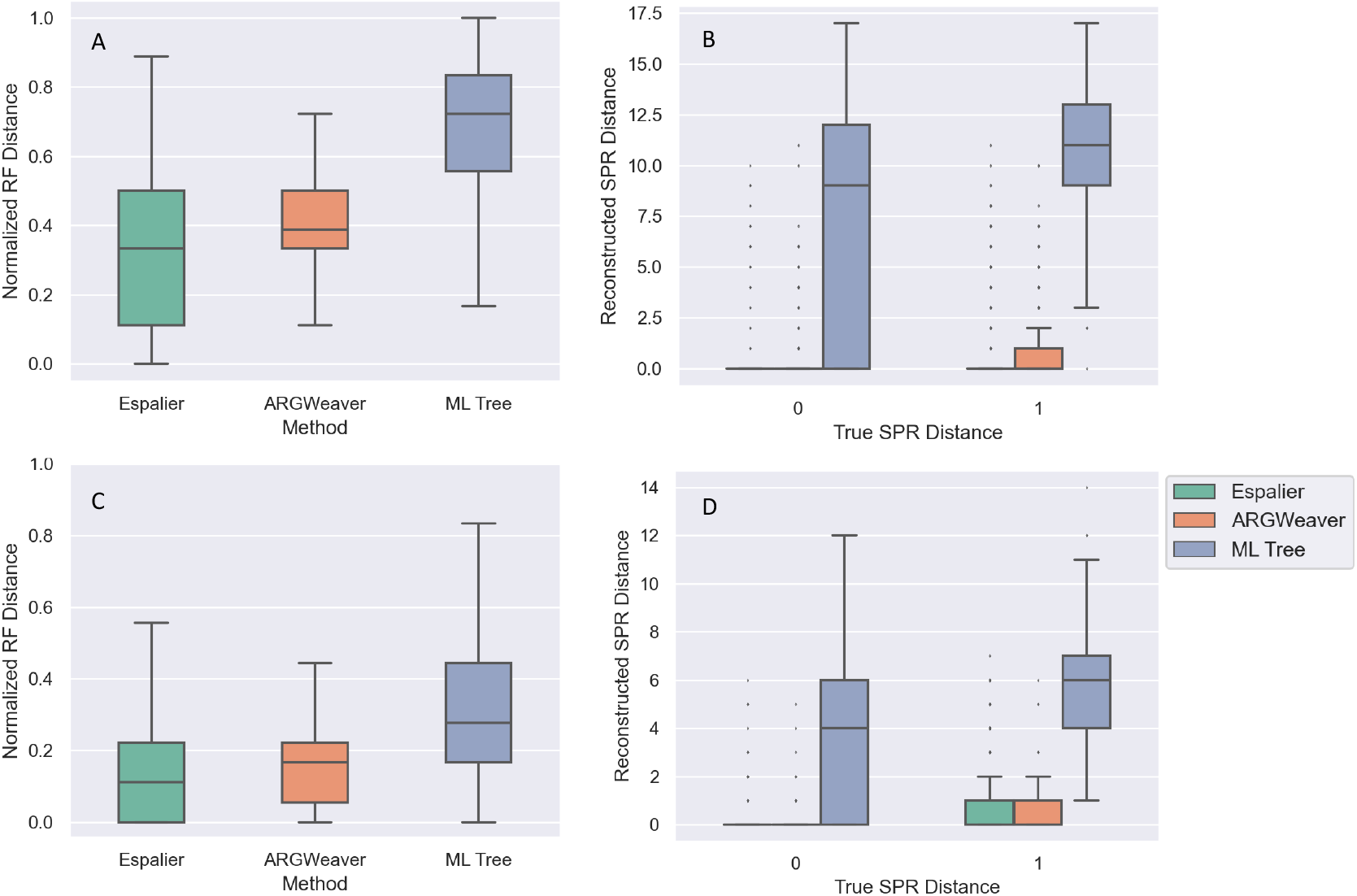
Performance reconstructing simulated ARGs. ARGs were reconstructed using either Espalier or ARGweaver and then local trees in the ARG were compared against the true local tree for each genomic region in simulated ARGs. For comparison, we also consider the maximum likelihood (ML) tree reconstructed by RAxML from sequence data for each genomic region individually. Simulations were performed at either low (A-B; *μ* = 0.01) or high (C-D; *μ* = 0.1) mutation rates. In **(A & C)**, normalized Robinson-Fould (RF) distances were computed between reconstructed and true local trees in the ARG. In **(C & D)**, the subtree-prune-regraft (SPR) distance between neighboring local trees reconstructed by each method is compared. The true SPR distance between neighboring local trees is always either zero or one depending on whether or not a recombination event has caused a topological rearrangement between trees. The sample size in all simulations was 20 with a genome length *L* = 1,000 base pairs

Neighboring local trees in the reconstructed ARGs are also quite close to one another in terms of SPR distances (Figure 6B). In the true ARGs, a single recombination event occurs at each recombination breakpoint between neighboring local trees, but recombination may or may not alter tree topology, such that the true SPR distance between local trees is always zero or one. When the true SPR distance is zero, the average SPR distance between neighboring trees is 0.24 for Espalier and 0.17 for ARGweaver, suggesting that errors in reconstructing local trees do not result in excessive discordance nor an excess of incorrectly imputed recombination events. When the true SPR distance is one, the average SPR distance between trees is 0.81 for ARGweaver but only 0.41 for Espalier. This suggests that at low mutation rates, Espalier may be overly conservative in allowing for topological changes between local trees and thus misses some true recombination events.

At higher mutation rates (*μ* = 0.1; *μ*/*r* = 100; π = 0.2), where there are more phylogenetically informative sites, local trees reconstructed by all methods (including the ML trees) are much closer to the true local trees. Espalier is slighly more accurate than ARGweaver in terms of reconstructing local tree topologies (Figure 6C) but both methods perform well in determining whether or not recombination has resulted in a topological change between neighboring local trees (Figure 6C). Overall, these results suggest that while Espalier may be less sensitive than ARGweaver in detecting recombination events when phylogenetic information is limiting, both methods perform well at “smoothing” differences between local trees by eliminating topological changes that are not well supported by sequence data and can accurately reconstruct ARGs when enough sequence diversity is present.

### Inferring recombination rates using the EM algorithm

Espalier allows demographic parameters to be estimated from ARGs while simultaneously reconstructing the ARG using the EM algorithm. Here we focus on estimating the recombination rate *r* under the coalescent with recombination since *r* is typically unknown but required for ARG inference. To test the performance of the EM algorithm in estimating *r*, we simulated ARGs together with mock sequence data and then obtained maximum likelihood estimates for the recombination rate, the number of recombination events, and the ARG.

Estimated recombination rates are strongly positively correlated with the true rates (Pearson correlation *R* = 0.51) but the accuracy of estimates varies widely across simulations and on averages rates are underestimated by about 10.7% (Figure 7A). The number of recombination events in the reconstructed ARGs is also slightly underestimated (Figure 7B). Error in the estimated number of recombination events is in turn strongly positively correlated with error in the estimated recombination rates (Figure 7C), suggesting that most of the error in estimated rates is attributable to a difficulty in correctly identifying recombination events. That both the rate and number of recombination events are underestimated is not surprising, as a large fraction of the recombination events (37% of events in our simulations) will not be detectable from sequence data due to the event having no effect on tree topology or the event occurring between genetically identical sequences. Overall, while inferring recombination rates appears inherently difficult, estimated rates are generally within a factor of two of their true values, suggesting that the EM algorithm is suitable for quickly obtaining ballpark estimates of recombination rates.

**Figure 7.**
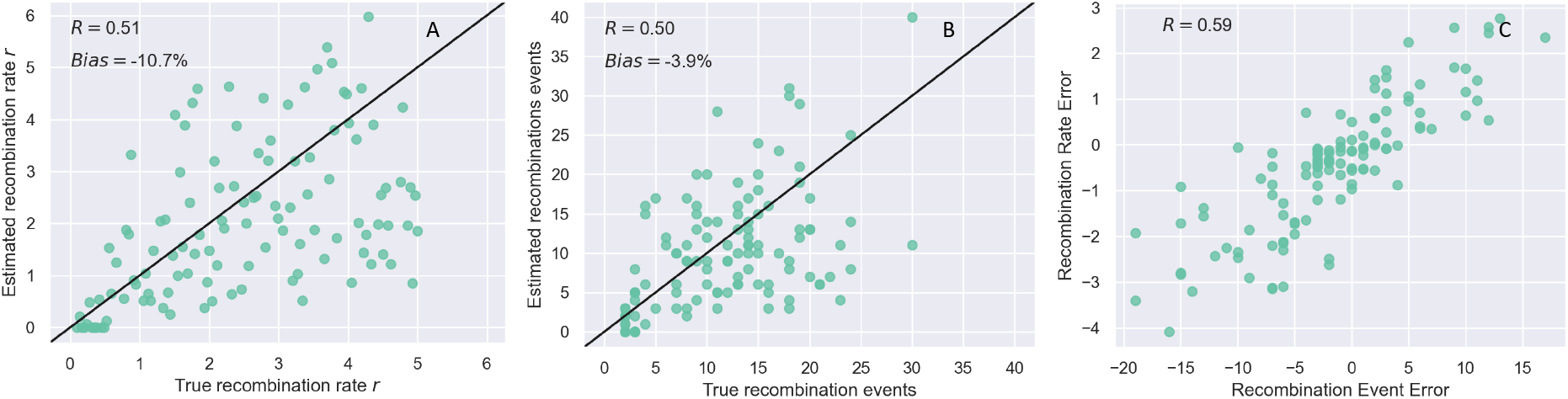
Performance of the EM algorithm in estimating recombination rates. True ARGs were simulated with recombination rates ranging from 1X (*r* = 0.1) to 50X (*r* = 5.0) times the assumed mutation rate (*μ* = 0.1; π = 2*N_e_μ* = 0.2). **(A)** True recombination rates versus maximum likelihood estimates obtained from the EM algorithm. R is the Pearson correlation coefficient between true and estimated values and bias is reported as the average error relative to the true value used in each simulation. **(B)** True versus estimated number of recombination events in the ARGs. **(C)** Absolute error in the inferred recombination rate is positively correlated with the error in number of recombination events in the reconstructed ARG. The sample size in all simulated ARGs was 20 with a genome length *L* = 10,000 base pairs.

#### Disentangling potyvirus tanglegrams

Potyviruses are a members of the genus *Potyvirus* in the family *Potyviridae*, the largest family of plant-infecting RNA viruses and arguably one of the most agronomically important due to major crop pathogens such as potato virus Y [Moury et al., 2017, Wylie et al., 2017]. Since the Neolithic agricultural revolution, potyviruses have rapidly radiated and expanded their host range and geographic distribution alongside their host plants [Gibbs et al., 2008, Gibbs and Ohshima, 2010]. Due to this rapid diversification, there is considerable uncertainty surrounding the deep phylogenetic relationships among the potyvirus species. Indeed, using a tanglegram to visualize how species-level phylogenetic relationships vary across the potyvirus genome shows widespread phylogenetic discordance between ML trees reconstructed from different regions of the genome without detected recombination breakpoints (see Figure 8B). However, potyviruses like other positive stranded RNA viruses frequently recombine, although the extent of interspecific recombination events between species is debated [Tan et al., 2004, Gibbs et al., 2020]. It is therefore unclear to what extent the discordance visualized in the tanglegrams is attributable to true phylogenetic conflict arising from recombination versus phylogenetic error.

**Figure 8.**
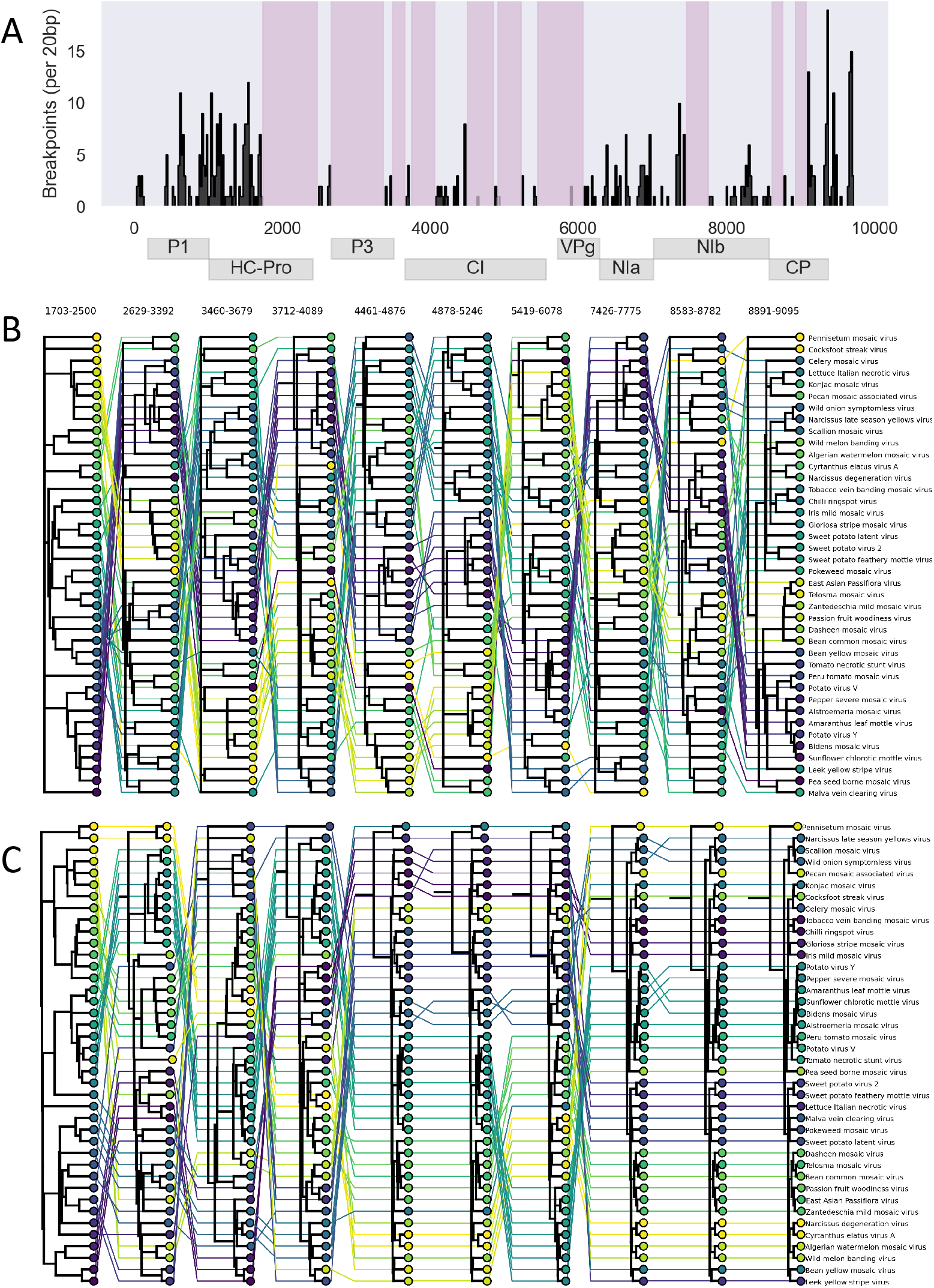
The recombinant phylogenetic history of the potyviruses. **(A)** The potyvirus genome consists of 10 protein coding regions translated as a single polyprotein. The histogram shows the number of potential recombination breakpoints per 20bp windows identified using 3SEQ. The 10 longest non-recombinant blocks (NRBs) chosen for phylogenetic analysis are shaded in magenta. **(B)** Tanglegram showing the discordant phylogenetic relationships of potyvirus species between ML trees reconstructed from each NRB individually. Horizontal lines connect the same species in each tree such that crossing lines indicate discordance. **(C)** A *disentangled* tanglegram for the local trees in the ARG reconstructed by Espalier. The tanglegrams shown here have been thinned to only include 40 of the 131 potyspecies species included in our full analysis.

In order to explore how much discordance between trees is attributable to recombination versus phylogenetic error, we reconstructed an ARG of the potyviruses using the EM algorithm in Espalier. The local trees within the reconstructed ARG produce a *disentangled* tanglegram in which discordances between local trees not strongly supported by the sequence data have been eliminated (Figure 8C). Note for purposes of visualization, only 40 potyvirus species are included in the tanglegrams shown in Figure 8. The ARG tanglegram shows dramatically less entanglement between trees suggesting that most of the discordance between local ML trees is likely attributable to phylogenetic error. While there remains considerable discordance between local trees in the ARG over the 5’ end of the genome, most of the imputed recombination events occur between lineages at the base of tree for which there is likely poor phylogenetic signal due to the combination of deep divergence times and rapid radiation. Thus, some of the putative interspecific recombination events identified in the ARG may still arise from phylogenetic error. However, local trees in the ARG over the 3’ end of the genome are largely concordant. For example, members of the potato virus Y clade including sunflower chlorotic mottle virus, bidens mosaic virus, amaranthus leaf mottle virus and pepper severe mosaic virus consistently cluster in a single clade in the six trees spanning the 3’ end of the genome. Overall, the potyviruses provide a clear example of how ARGs reconstructed by Espalier can be used to disentangle discordant relationships in order to provide a clearer picture of the evolutionary relationships between taxa.

## Discussion

Far from a single tree of life, genomic sequence data has revealed a far more complex “web” of life that is highly reticulate due to frequent recombination and horizontal transfers of genetic material [Smith et al., 1993, Soucy et al., 2015, Worobey and Holmes, 1999]. Indeed, outside of a few asexual or clonally evolving populations, the ancestry of most species can be thought of as a mosaic of phylogenetic histories that vary across the genome [Wiuf and Hein, 1999, Boni et al., 2007, Hubisz et al., 2020]. Extensive efforts have therefore been made to develop new phylogenetic methods accounting for recombination. Here, we demonstrate how maximum agreement forests, which have previously received little attention outside of computational phylogenetics [Hein et al., 1996, Rodrigues et al., 2007, Whidden and Zeh, 2009], can be used to build efficient algorithms for reconciling discordant phylogenetic trees and reconstructing ancestral recombination graphs.

Conceptually, our approach has much in common with other recent ARG reconstruction methods, but tries to strike a delicate balance between computational efficiency and the assumptions made by more approximate methods that limit their generalizibility. Like both ARGweaver [Rasmussen et al., 2014] and Arbores [Heine et al., 2018], Espalier formulates the ARG inference problem as a hidden Markov model with the goal of selecting a hidden path of trees across the genome under the assumptions of the sequential Markov coalescent [McVean and Cardin, 2005]. ARGweaver in particular appears to be very accurate at reconstructing ARGs but is also limited to small genomic datasets because it employs a computationally demanding MCMC algorithm to sample from the posterior distribution of ARGs. Arbores is similar to Espalier in that it uses SPR moves to search for compatible local trees in a tree path, but is likewise quite computationally expensive in that it considers all possible SPR moves between pairs of local trees, leading to a very large tree space to search. By contrast, the efficiency of Espalier lies in starting with pre-reconstructed local trees for each genomic region and then sampling a set of reconciled trees which are highly supported by the sequence data but have unnecessary topological discordances between trees removed. This greatly reduces the state space of the HMM in terms of the number of trees that need to be considered. However, unlike more approximate programs for inferring sequences of local trees across the genome [Kelleher et al., 2019, Speidel et al., 2019], Espalier makes additional advantage of MAFs to identify lineages that have likely recombined and then inserts the additional recombination events required to explain discordances between local trees.

Other than reconstructing initial local trees, most of the computational overhead of running Espalier lies in repeatedly computing MAFs between many pairs of trees. While running the MAF algorithm takes far less than a second even for trees with hundreds of taxa, repeatedly computing MAFs between many pairs of trees accumulates a large computational cost. Fortunately, we expect that the computational efficiency of the MAF algorithm can be greatly improved. For example, in order to choose which edge is cut in each iteration of the algorithm, the current algorithm copies tree objects in memory before making cuts. We therefore expect that operating on trees without copying tree objects will lead to drastic improvements in performance. Performance could be further improved using cluster-reduction algorithms that splits computation of the MAF into smaller sub-problems by clustering taxa into subsets [Linz and Semple, 2011, Whidden et al., 2014].

Recombination has also posed a significant challenge to fields such as phylogeography and phylodynamics, which seek to reconstruct evolutionary and demographic processes from reconstructed phylogenies. The ability to reconstruct ARGs in an efficient likelihood-based framework opens the way for performing demographic inference in a recombination-aware manner. In particular, a major advantage of computing ARGs is that they reveal which regions of the genome have unique ancestral histories and which regions provide redundant genealogical information. It may therefore be possible to more accurately and precisely infer demographic parameters like effective population sizes and migration rates from the multiple histories embedded within an ARG than any single gene tree or phylogeny [Li and Durbin, 2011, Müller et al., 2020].

One current limitation of Espalier for ARG reconstruction is that it does not infer recombination breakpoints and so breakpoints delimiting recombination-free regions of the genome need to be identified a priori. Nevertheless, the strategy we used to reconstruct potyvirus ARGs suggests a pragmatic and scalable computational pipeline for ARG inference. Breakpoints can first be identified using existing recombination detection software, recombination-free regions of the genome can be delimited between these breakpoints, local ML trees can be reconstructed from these recombinant-free regions, and then these local ML trees can be fed into Espalier for reconciliation. This basic strategy has been adopted for other viruses including coronaviruses [Boni et al., 2020, Jackson et al., 2021], and may work well for more highly recombining bacterial and eukaryotic organisms as long as there are regions of the genome where the recombination rate is low enough that multiple phylogenetically informative sites (e.g. SNPs) fall between recombination breakpoints.

Beyond reconstructing ARGs, the Espalier package provides a flexible toolkit for working with discordant phylogenetic histories using maximum agreement forests. The package includes methods and a command-line interface for computing MAFs, calculating SPR distances and tree reconciliation using iterative regrafting. The EM algorithm implemented in Espalier further allows for maximum-likelihood estimates of key demographic parameters like recombination rates to be rapidly inferred from ARGs. The package also allows for better visualization of discordant phylogenetic histories by removing unnecessary discordances, creating disentangled tangelgrams. Future work will allow for automated detection of recombination breakpoints and an expanded range of coalescent models for recombination-aware demographic inference.

## Acknowledgments

This research was supported by the USDA through FACT grant 2019-67021-29932. DAR was additionally supported through funding from USDA Hatch project 1016556.

## Supplemental Methods

### Iterative regrafting with prior information

In a situation where we are using the iterative regrafting algorithm to reconcile a local tree for a given genomic region against a reference tree, we can consider information about the local tree contained in the sequence data from sites neighboring but outside of the genomic region of interest. However, the phylogenetic history of sites outside of the region will likely increasingly differ with increasing distance from the region.

We therefore consider a prior *p*(*T_i_*|*S*_1:*L\i*_) on the local tree *T_i_* that considers information coming from sequence data outside of region *i*, denoted here as *S*_1:*L\i*_ where *S*_1:*L*\*i*_ ≔ *S*_1:*L*\[*l_i_*:*r_i_*]_. In particular, we exponentially discount information coming from sites outside of *i* based on their distance to *i*:

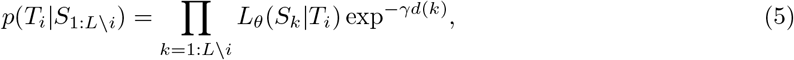

where *d*(*k*) is the genomic distance of site *k* from region *i* and *γ* determines the rate at which the contribution of sites outside of *i* decays with distance to *i*.

While including this prior adds another hyperparameter *γ* that needs to be tuned, knowing the recombination rate provides a natural way of parameterizing *γ*. We can set *γ* equal to the per site probability of a recombination event occuring anywhere in the ancestry of the sample, such that the prior information decays at a rate proportional to the the rate at which the probability that no recombination events has occurred between region *i* and site *k* decays. The per site probability of a recombination event is simply 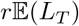, where *r* is the recombination rate per individual lineage and 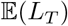 is the expected total length of the local tree. Under the standard neutral coalescent, the expected total length for a tree with *n* samples is:

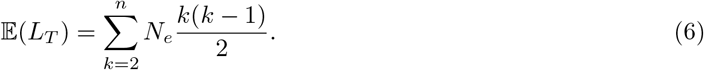

We can then use this prior to weight the likelihoods computed in the iterative regrafting algorithm by the likelihood of the sequence data outside of region *i* evolving under a given tree *T*:

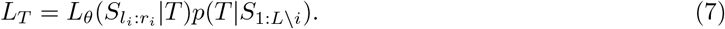

### A forward-backward algorithm for sampling tree paths

The Viterbi algorithm described in the main text finds the tree path with the most likely sequence of trees given the observed sequence data. Here, we describe a modified version of the forward-backward algorithm that allows us to randomly sample different tree paths from the joint distribution of trees *P*(*T*_1:*M*_|*S_I:M_*). In the forward pass, we compute the probabilities *f_i, k_* of being in each tree state *k* conditional on the sequence data observed up to region *i*, *P*(*T_i_*|*S*_1:*i*_). In the backward pass we then compute ”smoothed” probabilities that take into account the sequences for all regions. However, we modify the backwards pass of the algorithm to randomly sample a particular tree state at each time.

#### Algorithm 5: The forward-backward tree path sampler

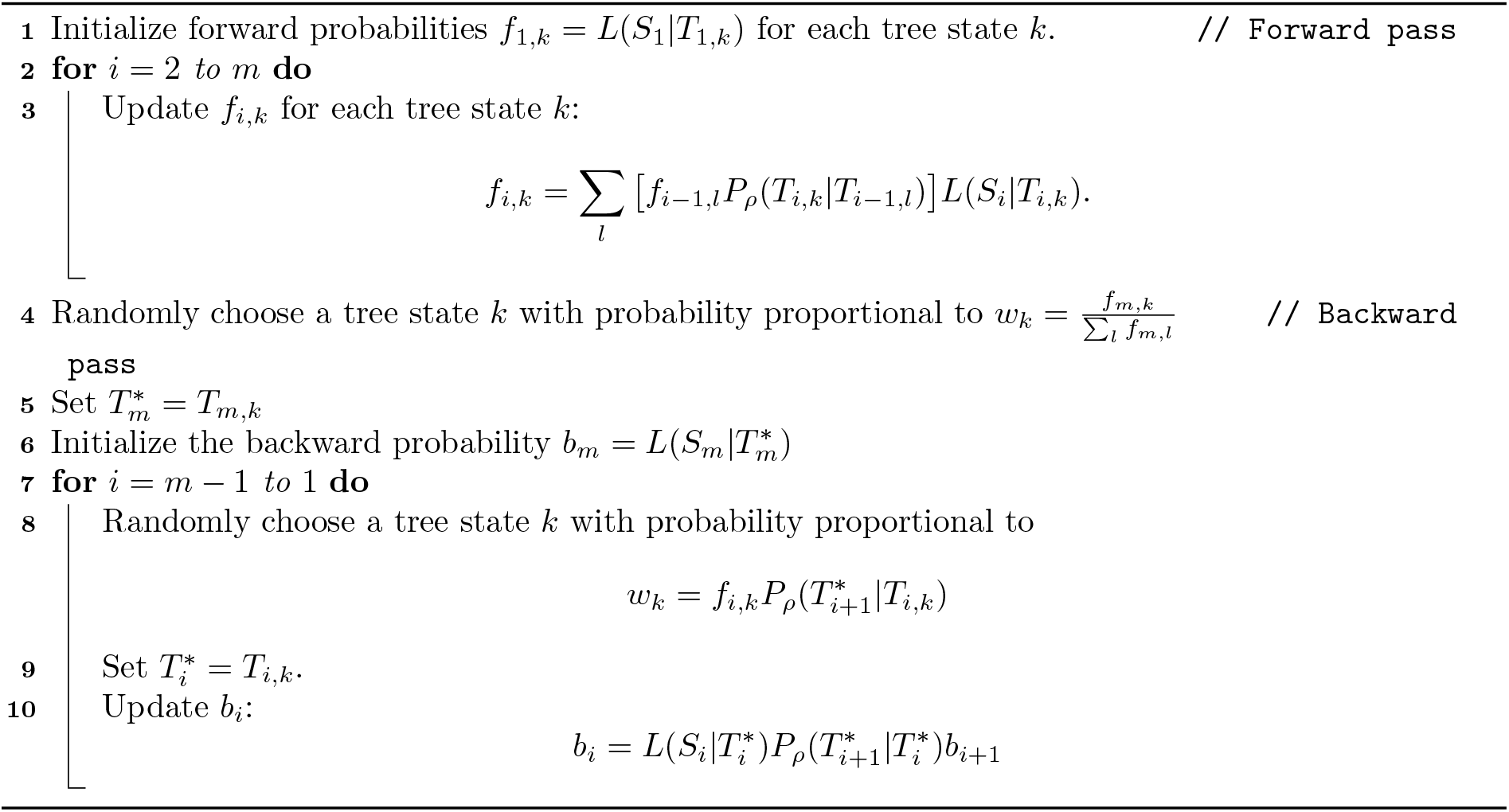

This algorithm differs from the standard forward-backward algorithm in that it couples computing the smoothed backward probabilities with a sampling step, such that we only compute the backward probability for the sampled state. Probabilistically sampling a tree state each step returns a tree path 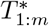 sampled from the joint posterior given in (1) and as a byproduct computes the likelihood of the sampled tree path through the backwards probabilities, as:

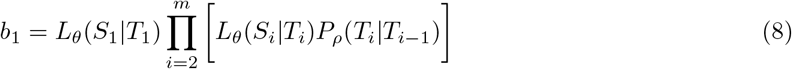

